# Physicochemical characterization, toxicity and *in vivo* biodistribution studies of a discoidal, lipid-based drug delivery vehicle: Lipodisq nanoparticles containing doxorubicin

**DOI:** 10.1101/2020.06.18.159087

**Authors:** Maria Lyngaas Torgersen, Peter J. Judge, Juan F. Bada Juarez, Abhilash D. Pandya, Markus Fusser, Charlie W. Davies, Matylda K. Maciejewska, Daniel J. Yin, Gunhild M. Mælandsmo, Tore Skotland, Anthony Watts, Kirsten Sandvig

## Abstract

Many promising pharmaceutically active compounds have low solubility in aqueous environments and their encapsulation into efficient drug delivery vehicles is crucial to increase their bioavailability. Lipodisq nanoparticles are approximately 10 nm in diameter and consist of a circular phospholipid bilayer, stabilized by an annulus of SMA (a hydrolysed copolymer of styrene and maleic anhydride). SMA is used extensively in structural biology to extract and stabilize integral membrane proteins for biophysical studies. Here, we assess the potential of these nanoparticles as drug delivery vehicles, determining their cytotoxicity and the *in vivo* excretion pathways of their polymer and lipid components. Doxorubicin-loaded Lipodisqs were cytotoxic across a panel of cancer cell lines, whereas nanoparticles without the drug had no effect on cell proliferation. Intracellular doxorubicin release from Lipodisqs in HeLa cells occurred in the low-pH environment of the endolysosomal system, consistent with the breakdown of the discoidal structure as the carboxylate groups of the SMA polymer become protonated. Biodistribution studies in mice showed that, unlike other nanoparticles injected intravenously, most of the Lipodisq components were recovered in the colon, consistent with rapid uptake by hepatocytes and excretion into bile. These data suggest that Lipodisqs have the potential to act as delivery vehicles for drugs and contrast agents.

## Introduction

Overcoming the low solubility of novel therapeutic compounds is a key challenge for the pharmaceutical industry [1]. Encapsulation of therapeutic agents within nanoparticles (NPs) may protect drug molecules from enzymatic degradation, maintain their efficacy and improve pharmacokinetics *in vivo* [2, 3]. NPs used for drug delivery typically have a diameter larger than 50 nm, and it is not known whether smaller NPs would give a better uptake in tumors due to the Enhanced Permeability and Retention (EPR) effect [4].

Lipid-based drug delivery systems, such as liposomes, micelles or nanodiscs [5, 6] are of interest since lipids can easily be metabolized in the body. They provide a unique amphipathic environment, with the ability to carry insoluble (or sparingly soluble) molecules in aqueous solutions. Doxil^®^/Caelyx^®^, a liposomal formulation containing doxorubicin (DOX) for cancer therapy, was FDA-approved in 1995, and was the first liposome-based product to enter the market [7, 8]. Several liposome-based products are now available [9]. Here we evaluate the ability of Lipodisq NPs (10 nm in diameter, 4.5 nm thickness) composed of a hydrolysed co-polymer of styrene and maleic anhydride (SMA) [10] and 1,2-dimyristoyl-*sn*-glycero-3-phosphocholine (DMPC; PC 14:0/14:0) [11, 12], to act as drug delivery vehicles for DOX. Since the Lipodisq nanoparticles are much smaller (10 – 20 nm) than the FDA approved liposomes (>50 nm), it is possible that Lipodisq NPs will give a higher uptake of drugs in tumors and better penetration into the tumor tissue. Additionally, liposomal larger NPs have limited stability in the body, and unknown long-term toxicity and poor EPR effect [13, 14].

DOX is active against a broad spectrum of tumors including breast, lung, gastric, ovarian, pancreatic, prostate, multiple myeloma and some types of leukaemia [15, 16]. The drug exerts its cytotoxicity by interchelating with DNA base pairs, as well as by interacting with several molecular targets such as DNA topoisomerase II [16]. DOX has low solubility in water, rendering its administration to patients difficult. Several side effects are observed in patients treated with DOX, such as nausea, skin irritation and fever, the most dramatic being heart failure (cardiomyopathy) [15, 17]. SMA polymer has been used extensively in structural biology to extract proteins directly from native membranes without the need for conventional detergents [10, 11]. The resulting Lipodisq NPs (also known as SMALPs), consisting of a coin-shaped lipid bilayer stabilised by a polymer annulus [11] are used extensively for characterisation of integral membrane proteins by different techniques, including cryo electron microscopy and X-ray crystallography [10]. Hydrophobic interactions drive the spontaneous self-assembly of Lipodisq NPs in aqueous conditions, creating a monodisperse suspension at physiological pH, temperature and ionic strength [10, 18].

Here we evaluate the ability of Lipodisq NPs composed of a hydrolysed co-polymer of styrene and maleic anhydride (SMA) [10] and 1,2-dimyristoyl-*sn*-glycero-3-phosphocholine (DMPC; PC 14:0/14:0) [11, 12], to act as drug delivery vehicles for DOX. The uptake of the NPs by cancer cells is studied, as well as the release of DOX in the acidic environments of lysosomes and the resulting cytotoxic effects across a panel of cancer cell lines. The *in vivo* fates of the NP components and the drug were followed after intravenous injection into mice and we observed uptake of the Lipodisq NPs by the liver, followed by rapid excretion in the bile. Our data suggest that intravenously injected Lipodisq NPs have a potential as delivery vehicles for drugs or as carriers for contrast agents, especially to the liver and lower gastrointestinal tract.

## Materials and Methods

### Reagents

Lipids including 1,2-dimyristoyl-*sn*-glycero-3-phosphocholine (DMPC), 1,2-dimyristoyl-*sn*-glycero-3-phospho-(1’-rac-glycerol) (DMPG) and 1,2-dilauroyl-*sn*-glycero-3-phospho-ethanolamine (DLPE), were purchased from Avanti Polar Lipids (Alabaster, AL). SMAnh (a non-hydrolyzed co-polymer of styrene:maleic anhydride in a 3:1 molar ratio) was kindly provided by Malvern Cosmeceutics Ltd (Worcester, UK). Doxorubicin (DOX) was from Cambridge Bioscience Ltd (Cambridge, UK) and Sulfo-Cyanine5 NHS ester (Cy5) were from Lumiprobe (Hannover, Germany). *N,N*-dimetlivlmetlianainide (DMF), Dimethyl sulfoxide (DMSO), Tris(2-carboxyethyl)phosphine (TCEP) and Concanamycin A (ConA) were obtained from Sigma-Aldrich (St. Louis, MO). Bafilomycin A1 (BafA1) was from Enzo Life Sciences, Farmingdale, NY.

### Preparation of Lipodisq batches

#### Preparation of Lipodisq NPs containing DOX

To prepare Lipodisq NPs containing different amounts of DOX, lipid (as a dry powder) and doxorubicin (fresh stock solution in ddH_2_O, added to yield 1, 5, or 10 % molar ratio) were dissolved in a chloroform:methanol (2:1 v/v) mixture, which was evaporated under a N_2_ stream, and the resulting lipid film was allowed to dry under vacuum (<10^-5^ Torr) overnight. Depending on the application, the film was rehydrated with either 50 mM phosphate buffer at pH 7.0 or 50 mM phosphate-buffered saline (PBS) at pH 7.4 (containing 137 mM NaCl and 2.7 mM KCl). The lipid suspension was then sonicated using a bath sonicator (5 min at room temperature, RT). Afterwards, 5-10 freeze thaw cycles were applied (in liquid nitrogen and then a water bath at 42 °C). Extrusion was performed using a Mini-Extruder (Avanti Polar Lipids, Alabaster, AL) and two 1000 μl syringes. The solution was extruded first using a Whatman 0.4 μm filter and then a Whatman 0.1 μm filter (10 mm diameter) (Sigma) at a temperature at least 5 °C higher than the phase transition temperature of the lipid when hydrated (approximately 30 °C for DMPC). Between 10 and 20 passages of the mixture through each type of filter were necessary to obtain a clear red suspension. Finally, SMA polymer was added in a 1:1.25 w/w lipid-to-SMA ratio. The mixture was then vortexed for a few seconds and maintained at 42 °C for 1 h, before the sample was centrifuged (14,000*g*, 5 min, RT).

The supernatant (of Lipodisq NPs containing DMPC and DOX) was concentrated prior to size exclusion chromatography using a Vivaspin column (MWCO 100 kDa, Millipore (Burlington, MA)) and then applied to a 5 ml HiTrap column (GE Healthcare, Chicago, IL) that had been equilibrated with phosphate or PBS buffer (depending on the application). Coloured fractions were collected, concentrated and stored at 4 °C for further experiments. An overview of all Lipodisq NP preparations is shown in Supplementary Table 1.

#### Preparation of a thiol derivative of SMA polymer (SMA-SH)

Non-hydrolysed styrene and maleic anhydride (SMAnh) polymer was conjugated to cysteamine using a method adapted from Lindhoud *et al*. [19]. Cysteamine (15 mg) was added to 1 g SMAnh and mixed in DMF. A volume of 700 μl of trimethylamine was added and the mixture stirred (30 min, RT). The resulting thiol derivative, SMAnh-SH, was precipitated by addition of approximately 30 ml of 1 M acetic acid and the precipitate was recovered after centrifugation (4000*g*, 5-10 min, RT). DMF was removed by washing with water (between 5 and 10 times) and the sample was then lyophilized. The polymer was hydrolysed to obtain SMA-SH (from SMAnh-SH) by dissolving SMAnh-SH powder in 0.1 M borate buffer, 100 mM dithiothreitol (DTT) pH 8.5 (100 °C, 4 h). After cooling, the solution was dialysed overnight (MWCO 3.5 kDa, 30 ml cassette from ThermoFisher, (Waltham, MA)) against 1 mM DTT, pH 8.0. The SMA-SH polymer was finally lyophilized and re-suspended in either PBS for further dye conjugation or in 1 mM DTT, pH 8.0 for longterm storage at a SMA-SH concentration of 125 mg/ml.

#### Preparation of fluorophore-labelled SMA (SMA-Cy5)

Sulfo-Cyanine5 NHS ester (Cy5) (Hannover, Germany) was dissolved in 20 μl DMSO and added directly to the SMA-SH previously dissolved in 50 mM phosphate buffer, pH 7.4 supplemented with 5 mM TCEP. The mixture was left to stir for 2 h in the dark at RT. Afterwards the polymer was extensively dialysed against a 50 mM phosphate buffer containing 1 mM DTT at pH 8.0. Unreacted dye and SMA-SH were removed by thin layer chromatography (TLC) using silica gel plates (Merck, Darmstadt, Germany) and a mobile phase of chloroform/methanol/water (5:5:1 v/v/v). SMA-Cy5 migrates with retention factors (Rf) between 0.4-0.7 and the free dye with an Rf=0.9. SMA-Cy5 was extracted from the silica by solubilisation in methanol, followed by centrifugation (4000*g*, 5 min, RT) and the supernatant was recovered and dried using a rotary evaporator. Once dried, the dye-labelled polymers were solubilized in PBS supplemented with 1 mM DTT, pH 8.0.

#### Preparation of fluorophore-labelled DLPE lipid (DLPE-Cy5)

Lipodisq NPs were also labelled with Cy5 by coupling to 1,2-dilauroyl-sn-glycero-3-phosphoethanolamine (DLPE; PE12:0/12:0). DLPE (4 mg, 6.90 μmol) and Sulfo-Cyanine5 NHS Ester (an *N*-hydroxysuccinimide ester derivative of the Cy5 dye Lumiprobe, Hannover, Germany) was conjugated to the lipid (Supplementary Figure 1) according to the literature [20], and purified by preparative TLC using a mobile phase of chloroform/methanol/water (95:25:4 v/v/v) (Supplementary Figure 2). The top band was recovered and analyzed by NMR and mass spectrometry to confirm that a single species had been purified (Supplementary Figure 3). This procedure resulted in formation of DLPE-Cy5 (6.78 mg, yield 93%), a bright blue viscous solid.

### Physicochemical characterization of Lipodisqs

The fluorescence spectrum of DOX is sensitive to the polarity of its microenvironment and this property was therefore used to investigate how DOX interacts with Lipodisq NPs. Furthermore, iodide (I^-^) quenches the fluorescence of DOX in solution [21, 22]. For the fluorescence quenching experiments, two samples were required for each lipid preparation: one containing KI (100 mM) and the other containing KCl (100 mM) in phosphate buffer (pH 7.0). The buffer was added at the beginning of the preparation of the Lipodisq suspensions (addition of stock solution of 1 M KCl or KI to the Lipodisq-Dox sample). The fluorescence measurements were performed using a LS50B Luminescence Spectrometer from Perkin Elmer (Seer Green, UK). Excitation was performed at 480 nm and the emission spectra were recorded from 500 nm to 650 nm. These measurements were performed in 500 μl cuvettes with samples diluted 10 times with phosphate buffer. The temperature was maintained at 20 °C using a water bath.

Lipodisq particle size in each of the preparations was measured by dynamic light scattering (DLS) using a Malvern Zetasizer Nano S instrument (633 nm; disposable cuvettes), and data were processed using Malvern Zetasizer software (Great Malvern, UK).

^1^H and ^13^C NMR spectra were recorded using a Bruker 500 MHz spectrometer running Topspin™ software, measured against a residual solvent peak as an internal standard. Chemical shifts (δ) are given in parts per million (ppm). The ^1^H NMR spectra are reported as follows: δ/ppm (assignment, multiplicity, number of protons). Multiplicity is abbreviated as follows: s = singlet, m = multiplet. Compound names are those generated by ChemBioDraw™ (CambridgeSoft) following IUPAC nomenclature. The NMR assignment numbering used is arbitrary and does not follow any particular convention. The ^13^C NMR spectra are reported in δ/ppm. Two-dimensional (COSY, HSQC, HMBC) NMR spectroscopy was used to assist the assignment of signals in the ^1^H and ^13^C NMR spectra. All NMR experiments were carried out at 25 °C unless otherwise stated.

High-resolution mass spectra were recorded on a Bruker ESI MicroTof mass spectrometer (Chemistry Research Laboratory, University of Oxford, UK).

### Cell lines and treatments

All cancer cell lines used in this study were obtained from ATCC and routinely tested for mycoplasma. The MDA-MB-231 (triple-negative breast cancer, Claudin low) and SW480 (colorectal adenocarcinoma) were cultured in RPMI; the MDA-MB-468 (triple negative breast cancer, basal), HeLa (cervical cancer), HCT116 (colorectal cancer), SKBR3 (breast cancer, HER2+), and MCF7 (breast cancer, luminal A) were cultured in DMEM. All media were fortified with 10% (v/v) fetal calf serum (Sigma-Aldrich, St. Louis, MO) and 100 units/ml penicillin/streptomycin (PenStrep^®^, Sigma-Aldrich, St. Louis, MO). The cells were seeded one or two days prior to experiments. For inhibitor studies, cells were pre-treated with the inhibitors at the indicated concentration for 1 h before addition of Lipodisq NPs.

### Cell proliferation measured by [^3^H]thymidine incorporation

Incorporation of [^3^H]thymidine into DNA was used to estimate cell proliferation. Cells seeded at an initial cell density of 40,000-60,000 per well in a 24 well plate were treated with either free DOX, empty Lipodisqs (LQ-E), or Lipodisqs containing 1% DOX (LQ-1%), 5% DOX (LQ-5%) or 10% DOX (LQ-10%), where the percentage of DOX indicates the molar ratio to the lipid. The compositions of all Lipodisq batches used are listed in Supplementary Table 1. After 6-24 h at 37 °C (as specified in each figure legend) the cell medium was aspirated and substituted with serum-free cell medium containing [^3^H]thymidine (2 μCi/ml, Perkin Elmer, Waltham, MA). The incubation was continued for 30 min at 37 °C. The medium was removed and 5% (w/v) trichloroacetic acid (TCA) was added. After 5 min the cells were washed twice with TCA and solubilized with 200 μl of 0.1M KOH, before mixing with 3 ml scintillation fluid (Perkin Elmer, Waltham, MA). The radioactivity was counted for 1 min in a scintillation counter (Tri-Carb 2100TR, Packard Bioscience, Arvada, CO). To assess the stability of the Lipodisqs and the release of DOX from the Lipodisqs, the Lipodisqs were pre-treated as described in each figure legend. Thereafter, the treated Lipodisq solution was passed through a Centrifree^®^ Ultrafiltration Device (Sigma-Aldrich, St. Louis, MO) that is commonly used for separating free from bound therapeutic drugs in biological samples. The filtration was performed in a Universal 320 R centrifuge (Hettich, Tuttlingen, Germany) at 1800*g* for 7 min.

### MTT cell viability assay

Cytotoxicity was assessed by the MTT (4,5-dimethylthiazol-2-yl)-2,5-diphenyltetrazolium bromide) cell viability test. Cells seeded at an initial density of 5000 or 8000 cells per well in a 96 well plate were treated with increasing concentrations of free DOX or with equivalent concentrations of DOX-loaded Lipodisqs (LQ-10%) and the same dilutions of empty Lipodisqs (LQ-E). After 48 h at 37 °C the cell medium was aspirated, 100 μl of medium containing a final concentration of 250 μg MTT/ml were added, and the incubation was continued for 3 h at 37 °C. The formazan-particles formed were subsequently dissolved in DMSO/0.25% NH_4_Cl during vigorous shaking for 10 min, and the absorbance was read in a plate reader (BioTek, Winooski, VT) at 570 nm (subtracting background absorbance at 650 nm).

### Live-cell confocal fluorescence microscopy

HeLa cells were seeded in 8-well Lab-Tek chamber slides (VWR, Radnor, PA) at a density of 30,000 cells per well one day prior to experiments. For visualisation of lysosomes, the cells were transduced with the lysosomal marker BacMam CellLight™ LysosomesRFP (Invitrogen, Carlsbad, CA) according to the manufacturer’s recommendations. Empty Cy5-labeled Lipodisqs (LQ-E) were added in fresh complete medium 16 h later. Lysosomes were also detected with the acid-sensitive lysosomal dye Lysotracker Green (100 nM, Invitrogen, Carlsbad, CA). The dye was added 5 min before imaging. The cells were imaged using a Zeiss LSM780 laser scanning confocal microscope (Carl Zeiss Microscopy, Jena, Germany) equipped with an Ar-Laser multiline (458/488/514 nm), a DPSS-561 10 laser (561 nm), and a Zeiss Plan-Apochromat 63×/1.40 Oil DIC M27 objective. For visualisation of DOX, fluorescence in DOX-loaded Lipodisqs LQ-10% was used.

### Biodistribution in mice

The animal experiments were approved and performed according to the Norwegian Animal Research Authority (Permit number 7837) and were conducted according to the regulations of the Federation of European Laboratory Animal Science Associations (FELASA). The mice were kept under pathogen-free conditions, at constant temperature (21.5 ± 0.5 °C) and humidity (55 ± 5%); 15 air changes/h and a 12 h light/dark cycle. They had access to distilled water *ad libitum*. All mice used were female athymic nude foxn1^nu^ mice (age 5-6 weeks and body weights of 18-20 g), locally bred at the Department of Comparative Medicine, Oslo University Hospital, Norway.

Lipodisqs labelled with Cy5 either bound to the polymer SMA or to the lipid DLPE (Supplementary Table 1) were used to study the biodistribution in healthy mice using an IVIS^®^ Spectrum *in vivo* imaging system (Perkin Elmer). Mice were intravenously injected with these two formulations of Lipodisqs (injected volume of 100 μl with a dose of 100 mg/kg lipid) and saline as a control. The mice were given 4% sevoflurane gas anaesthesia using multiple masks. The excitation/emission wavelength pair of 640/680 nm was found to give the best signal-to-noise ratio and this setting was thus used for imaging of the NPs. As described below, two studies were performed and whole body images were obtained either immediately after injection of substance or 4 and 24 h after injection; the animals were then sacrificed by cervical dislocation and organs were harvested at the time points described. The organs were imaged *ex vivo* with the IVIS scanner using the same settings as above. Relative signal intensity in the organs was calculated, using Living Image software (Perkin Elmer), as radiant efficiency (Emission light [photons/sec/cm^2^/str]/ Excitation light [μW/cm^2^] ×10^9^) per pixel of the region of interest, which was drawn around the respective organ.

## Results and discussion

### DOX is stably incorporated into 10 nm Lipodisqs

Lipodisq NPs containing DOX were formed by the addition of SMA polymer to liposome suspensions containing the drug. The self-assembly of the discoidal particles is spontaneous and is driven by hydrophobic interactions, as the styrene groups of the polymer interact directly with the non-polar lipid tails [10]. NP formation was verified by dynamic light scattering and all Lipodisq preparations containing DOX were found to have a similar size with a diameter of 10 ± 2 nm (data not shown) [11].

DOX has previously been shown to partition into model membranes containing DMPC, by allowing its planar aromatic rings to insert between the lipid acyl chains and its polar headgroup (containing a primary amine) to interact with the negatively charged lipid phosphate groups [23]. We proposed that DOX would be incorporated into the Lipodisq NPs in a similar manner and that the delocalised electron system, that gives rise to the fluorescence properties of the drug, would be inaccessible to the aqueous phase. The pKa of the primary amine group in DOX is 8.2 (PubChem database). In our experiments, conducted at pH 7.0, the amino carbohydrate moiety of DOX would be expected to be mostly positively charged in solution, thus preventing deep penetration into the lipid layer. In addition, the lateral pressure profile, also replicated by the Lipodisq NPs, has been reported to hinder the deep penetration of DOX into lipid bilayers [24]. The DOX fluorescence emission spectrum is highly sensitive to its microenvironment and the drug dimerizes at high concentrations in solution (K_d_ ~ 0.2 mM), resulting in self-quenching [25, 26]. Fluorescence quenching of the DOX may also be achieved with the addition of I^-^ ions, which can interact with the aromatic ring system (Figure 1A) [21, 22]. We therefore compared the relative fluorescence intensities of free DOX (in solution) and DOX in Lipodisqs (Supplementary Table 1) in the presence of 100 mM Cl^-^ and 100 mM I^-^ (Figure 1B). An increased fluorescence intensity of DOX in the Lipodisq NPs relative to the solution sample was observed; this was likely caused by a reduction in the dimerization of DOX upon incorporation into the NPs [25, 26]. DOX in Lipodisq NPs was quenched much less by iodide than DOX in free solution (Figure 1C), suggesting that in the NPs, the drug is not solvent accessible and is protected by the lipid environment of the Lipodisq. Lipodisqs comprised of DMPG instead of DMPC, showed very similar (but slightly reduced) quenching behaviour to DMPC Lipodisqs (Figure 1B-C), indicating that lipid charge does not markedly affect stability or formation of Lipodisqs – DMPG is anionic at pH>4.0.

**Figure 1.**
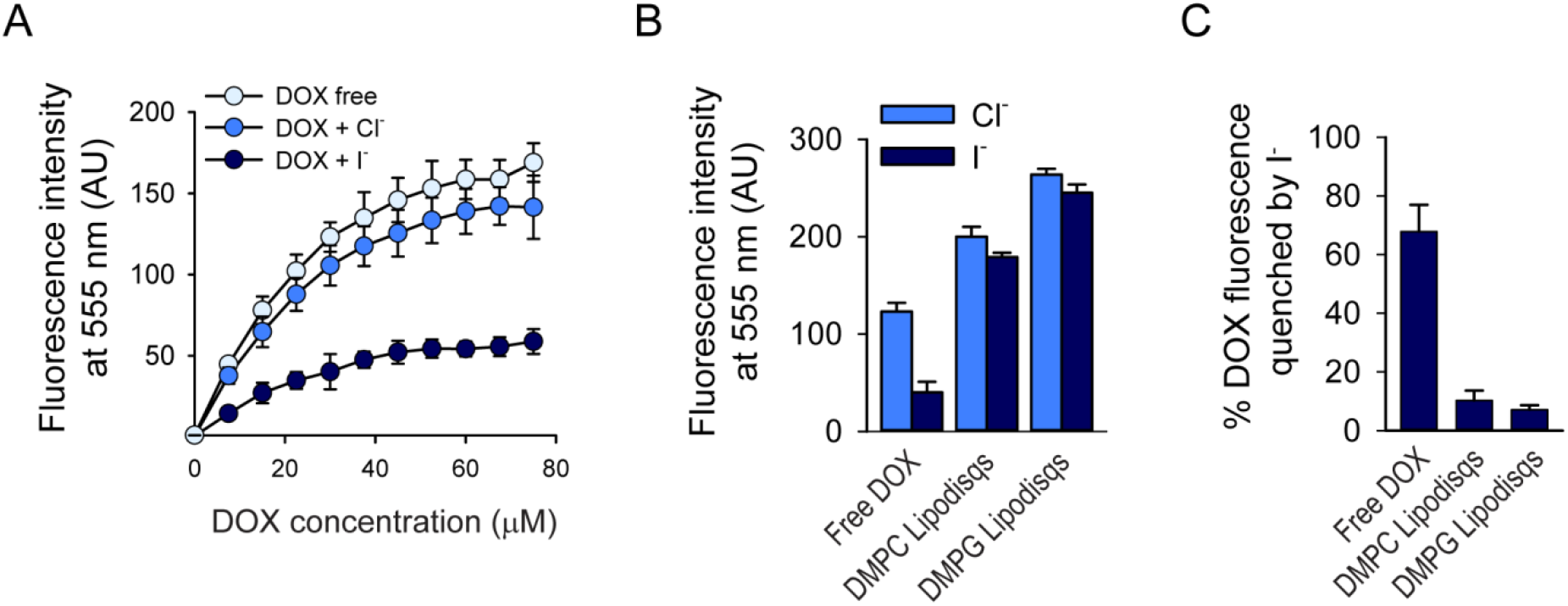
DOX is less susceptible to fluorescence quenching when incorporated in Lipodisqs than in solution. **A.** Fluorescence measured with increasing concentrations of DOX and the effect of adding 100 mM KI or KCl. **B-C**. Comparison of relative fluorescence intensities and quenching of DOX in solution vs in Lipodisqs. **B.** Peak fluorescence measurements of DOX formulations in 100 mM KCl vs 100 mM KI. **C.** Percentage of DOX fluorescence quenched by I^-^ against the Cl^-^ control. Experiments were performed with 30 μM DOX, corresponding to a 1% drug:lipid molar ratio in Lipodisqs containing 2 mg of DMPC or DMPG. All experiments were performed in 50 mM phosphate buffer, pH 7.0, n=5. Error bars show one standard deviation.

The sensitivity of the fluorescence peak ratios of DOX to the polarity of its environment [27] was used to probe the average local dielectric constant in the microenvironment of the drug. DOX was dissolved in different solvents (listed in Supplementary Table 2); the dielectric constants were taken from Wohlfarth [28]. The ratios of the intensities of the emission peaks, i.e. 555 nm/558 nm and 630 nm/588 nm, were plotted (Supplementary Figure 4). The 555 nm/558 nm ratio was 1.09 and the 630 nm/588 nm ratio was 0.21, indicating that the average local dielectric constant of DOX is between 50 and 110, consistent with the position of the drug close to the charged lipid headgroups and glycerol moiety [24]. Together these fluorescence data suggest that the location of DOX is likely to be around the glycerol moieties of the phospholipids, with the positive charge from the amino group of DOX protruding from the Lipodisq NPs (Figure 2).

**Figure 2.**
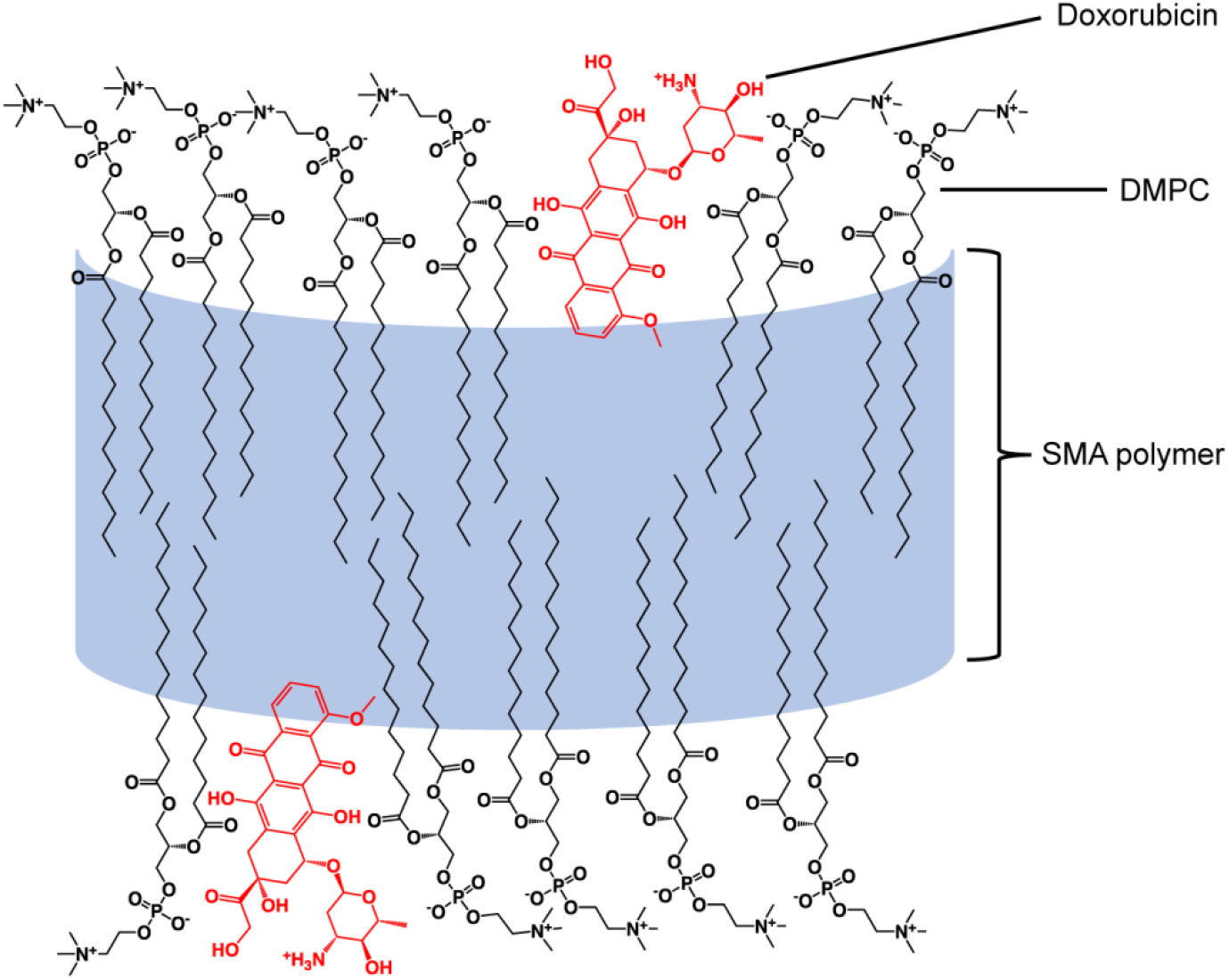
Illustration of the proposed location of DOX within a DMPC Lipodisq. The DOX molecules were adapted from Dupou-Cezanne et al. [24], whilst the DMPC molecules were from the Avanti website.

### DOX is rapidly released from Lipodisqs upon cellular uptake

To assess the cytotoxicity of DOX-loaded Lipodisq NPs, their effect on cell proliferation was determined by measuring thymidine incorporation. HeLa cells were treated for 24 h with increasing concentrations of either empty Lipodisqs (LQ-E), or Lipodisqs containing 1% DOX (LQ-1%), 5% DOX (LQ-5%) or 10% DOX (LQ-10%). The DOX-containing NPs clearly reduced cell proliferation (Figure 3A). When the data were plotted as a function of NP DOX concentration, the curves fully overlapped (Figure 3B), suggesting that Lipodisq NP cytotoxicity strictly correlates with DOX content.

**Figure 3.**
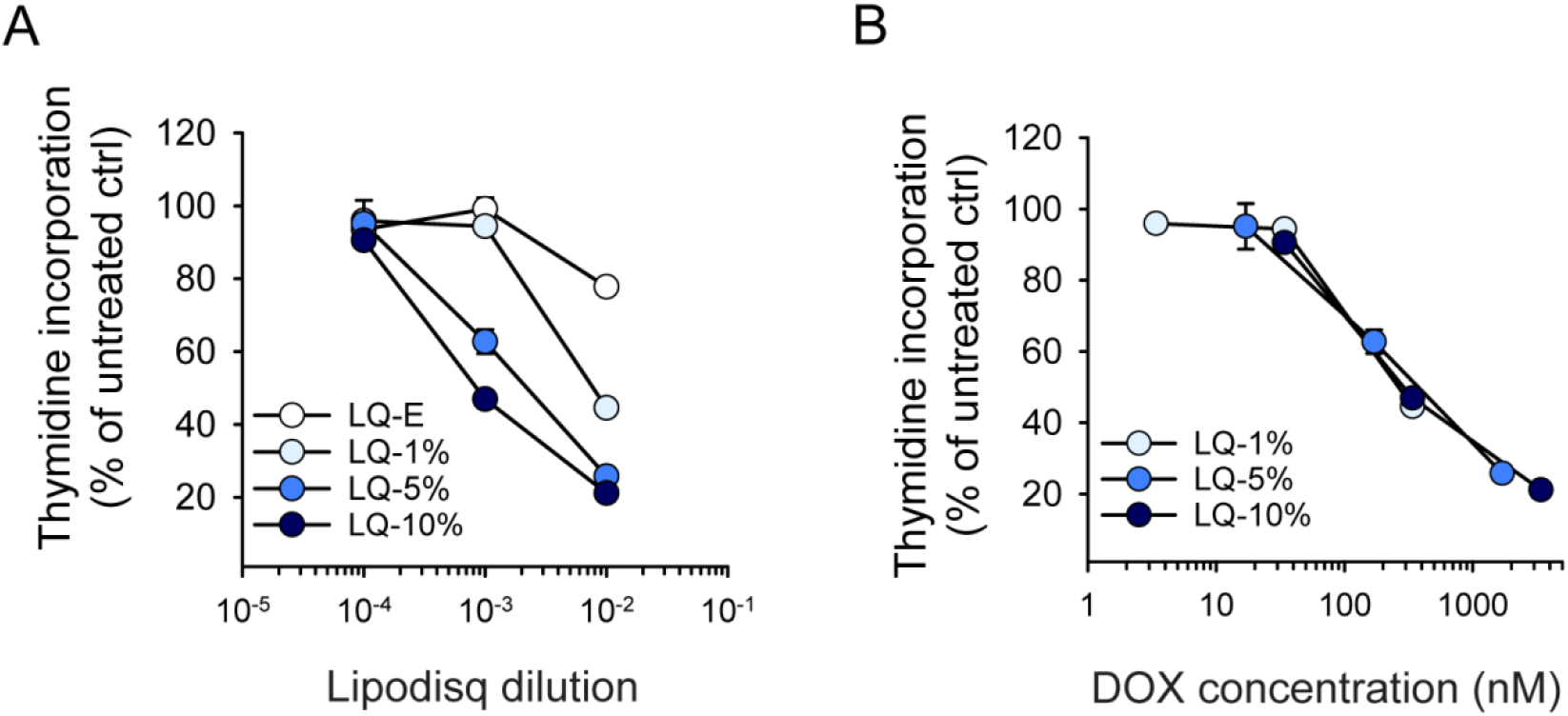
Cytotoxicity of Lipodisqs correlates with DOX loading. **A-B.** HeLa cells were treated with increasing dilutions of empty Lipodisqs (LQ-E) or Lipodisqs loaded with 1% (LQ-1%), 5% (LQ-5%), or 10% DOX (LQ-10%) for 16 h. Then thymidine incorporation was assessed by incubation with [^3^H]thymidine for 30 min, and the data were normalised to untreated cells. Thymidine incorporation was either plotted as a function of Lipodisq dilution (**A**) or DOX loading (**B**).

Thymidine incorporation into HeLa cells was used to determine how and when DOX is released from Lipodisq NPs. First, we assessed whether DOX is released from the NPs in the absence of cells, for example by non-specific interactions with serum proteins in complete growth medium (see schematic drawing of workflow in Supplementary Figure 5A). LQ-5% was incubated in complete medium for increasing periods of time (3-29 h) at 37 °C before filtration through a Centrifree spin column, to separate the free from the bound drug. Free DOX molecules pass through the column, whereas drug incorporated in NPs is retained by the Ultracel^®^ regenerated cellulose membrane. No reduction in cell proliferation was observed when cells were treated with the filtrate, suggesting that Lipodisq NPs are stable in complete medium (Supplementary Figure 5B). In contrast, unfiltered NPs displayed potent cytotoxicity, as expected (Supplementary Figure 5B). Free DOX displayed the same cytotoxicity with or without filtration (Supplementary Figure 5C), demonstrating that free drug molecules can pass freely through the column.

The possibility existed that DOX is released immediately upon contact between the Lipodisqs and the cellular surface. To test this, HeLa cells were pre-incubated with LQ-10% both for 30 min on ice and for increasing time at 37°C, before washing away unbound NPs, after which the incubation continued. When cell proliferation was assessed 6 h after initial NP addition, the data showed that potent release of DOX required at least 2-4 h of incubation at 37 °C (Supplementary Figure 6), suggesting that DOX release requires cellular internalization.

After 4 h of incubation with LQ-10%, a punctate, cytoplasmic signal from the Lipodisq-conjugated Cy5 was evident in HeLa cells (Figure 4A), consistent with the uptake of the NPs into the cell. The red fluorescence of DOX partially overlapped with the punctate Cy5 staining, but was also detectable in the nucleus, as expected from free DOX molecules, and consistent with the cytotoxicity observed after 4 h of incubation with Lipodisqs. The punctate Cy5-positive structures resembled lysosomes, and indeed, a substantial overlap between the Cy5 signal from LQ-10% and lysosomes stained by either CellLight LysosomesRFP or the lysosomal dye Lysotracker Green (Figure 4B-C) was observed. Together, these data suggest that Lipodisqs are transported to lysosomes for degradation and that DOX is released from within the endolysosomal system.

**Figure 4.**
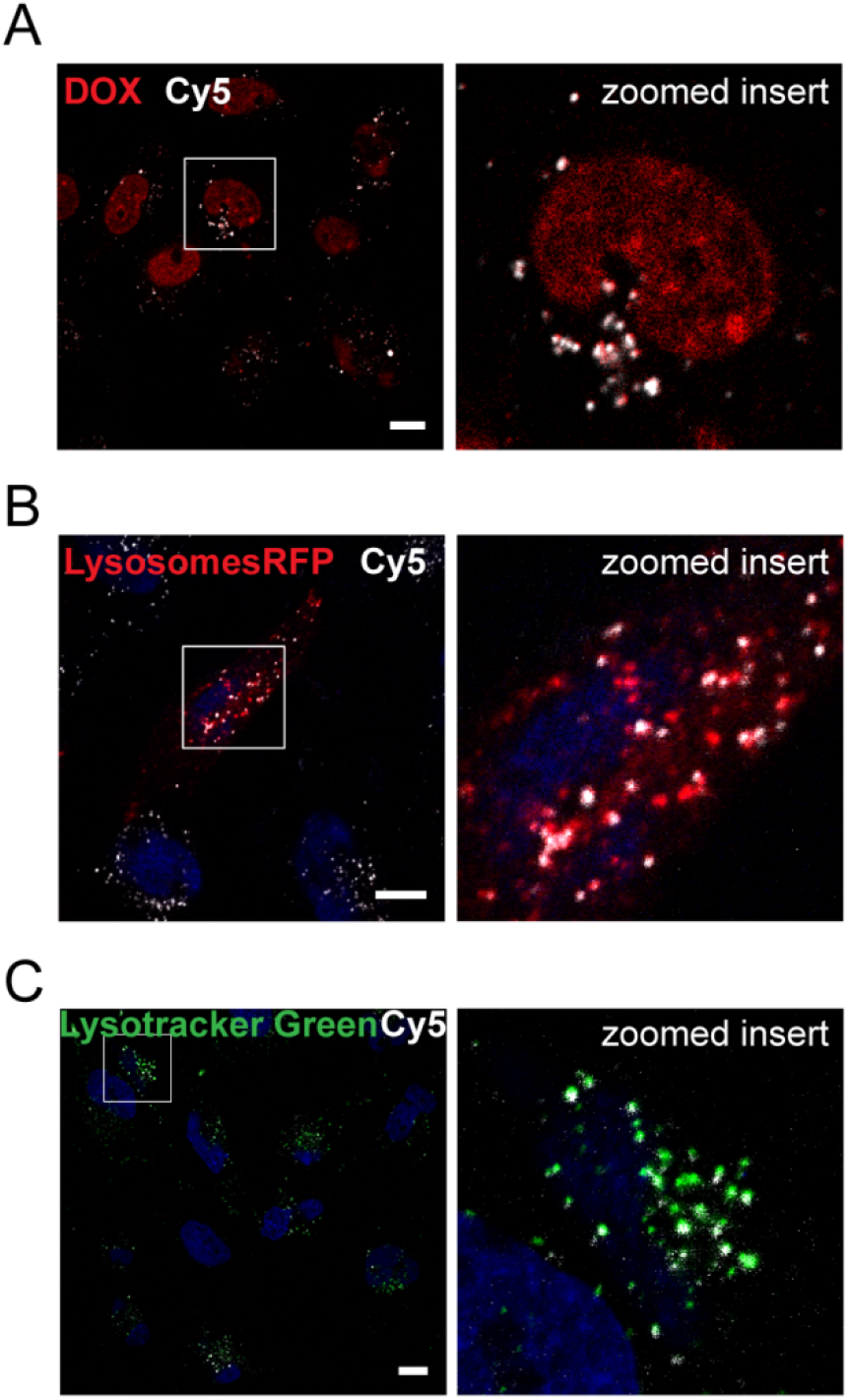
Lipodisqs are rapidly internalized to lysosomes in HeLa cells. **A.** HeLa cells were treated with Cy5-labeled Lipodisqs containing 10% DOX (LQ-10%, 1:100) for 4 h. **B-C:** For staining of lysosomes, HeLa cells were either transduced with BacMam CellLight™ LysosomesRFP and incubated for 16 h (**B**), or stained with Lysotracker Green (100 nM) 5 min before imaging (**C**). The cells were incubated with empty Cy5-labeled Lipodisqs (LQ-E, 1:10) for 4 h and the nuclear stain Hoechst (1.6 μM) was added before imaging. Images were acquired by an LSM780 confocal microscope in live cell mode. Scale bar; 10 μm. The right panels show enlargements of the indicated areas.

### DOX is released from Lipodisq NPs at low pH

The internal environment of lysosomes has a pH of 4.5-5.0, maintained by lysosomal vATPases [29]. To assess whether a low pH is required for DOX release, we incubated LQ-10% for 4 h in PBS pH 7.4 or 5.0 and compared the cytotoxic effect of the filtrate as a measure of DOX release. As shown in Figure 5A, incubation at pH 5.0 renders the filtrate significantly more cytotoxic than incubation at pH 7.4, suggesting that the release of DOX is increased at pH 5.0. Of note, treatment of free DOX at different pH values does not alter the cytotoxicity of the drug itself (Figure 5A). The release of DOX at low pH was not a spontaneous process, since at least 4 h of incubation at pH 5.0 was required to provide the filtrate with cytotoxic properties (Supplementary Figure 7). The release of DOX at pH 5.0 fits well with earlier data showing that the SMA polymer becomes protonated at this pH [10] and then dissociates from the lipid. We therefore propose that release of drug from the discoidal NPs occurs as a result of a change in the lipid morphology (and breakdown of the disc structure) at low pH [10]. We cannot exclude the possibility that also lipases that are active at low pH contribute to disc breakdown and DOX release.

**Figure 5.**
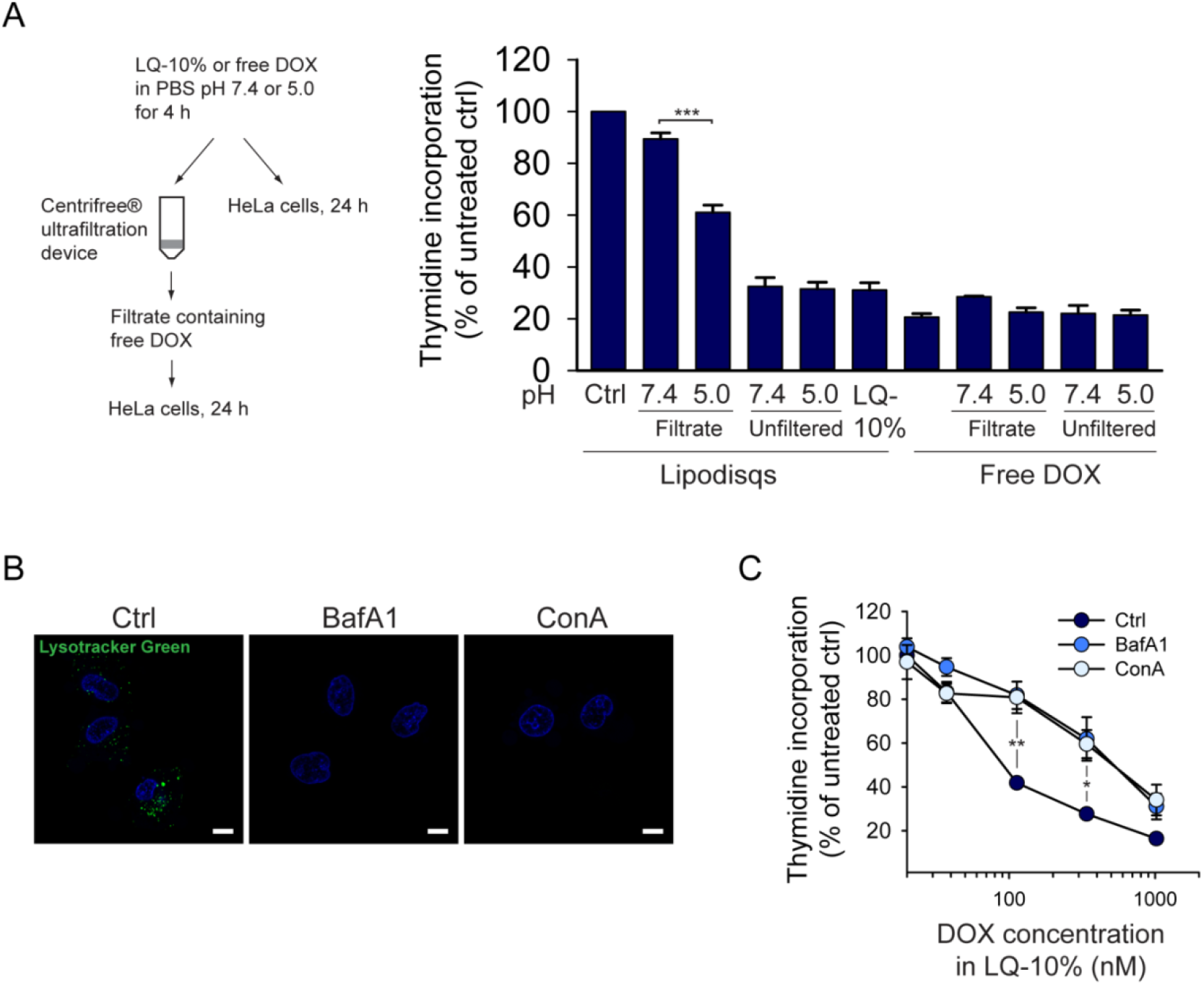
Low pH environment leads to release of DOX from Lipodisqs. **A.** LQ-10% (1:25) or free DOX (13.6 μM) was incubated in PBS pH 7.4 or 5.0 at 37 °C for 4 h. Then half of the solution was filtered through a Centrifree^®^ spin column to separate free DOX from Lipodisq-bound drug. Both the filtrate and the unfiltered half were diluted further (1:40) in complete medium and added to HeLa cells. Untreated LQ-10% was added as a control. The cells were incubated for 24 h before the level of thymidine incorporation was determined by incubation with [^3^H]thymidine for 30 min. The data were normalised to untreated control cells. **B.** HeLa cells were treated with the vATPase inhibitors Bafilomycin A1 (BafA1, 25 nM) or Concanamycin A (ConA, 25 nM) for 1 h, then the acid-sensitive lysosomal stain Lysotracker Green (100 nM) and the nuclear stain Hoechst (1.6 μM) were added and images were acquired after 5 min. Scale bar; 10 μm. **C.** HeLa cells were treated with increasing concentrations of LQ-10% for 24 h in the absence or presence of BafA1 (50 nM) or ConA (40 nM). Then thymidine incorporation was assessed, and the data were normalised to non Lipodisq-treated cells for each series. The graphs show mean values ± SEM from three independent experiments except for free DOX in A, where the mean values from two independent experiments are shown and the error bars show deviation from the mean. *, p < 0.05; **, p < 0.01; ***, p < 0.001.

To assess whether the degree of endosomal and lysosomal acidification affects the cytotoxicity of Lipodisq NPs, HeLa cells were treated with well-known inhibitors of lysosomal vATPases, Bafilomycin A1 (BafA1) and Concanamycin A (ConA) [30]. To verify that acidification was abolished by the inhibitors, HeLa cells were stained with the acidsensitive Lysotracker Green dye, which is trapped selectively within acidic compartments when protonated. In the presence of the two inhibitors, the punctate staining previously observed in control cells was completely abolished (Figure 5B) and the cytotoxicity of LQ-10% was significantly reduced (Figure 5C), consistent with a reduced release of DOX from the NPs.

### DOX-loaded Lipodisqs exert cytotoxic action across a panel of cancer cell lines

To test whether the data obtained in HeLa cells are representative for other *in vitro* systems, the cytotoxicity of empty Lipodisqs (LQ-E) with that of DOX-loaded Lipodisqs (LQ-10%) and free DOX was compared across a panel of cell lines from breast and colorectal cancer origin; MCF7, MDA-MB-468, SKBR3, and MDA-MB-231, or HCT116 and SW480, respectively. Although the cytotoxic effect differed between these cell lines, the toxicity of the DOX-loaded Lipodisqs was similar to that obtained with free DOX and much higher than that obtained with the empty Lipodisqs (Figure 6). Comparable data were obtained after 72 h of incubation (our unpublished observations). Moreover, LQ-10% exerted different cellular effects depending on the concentrations added. At relatively low concentrations (approx. 340 nM DOX content) typical signs of senescence were observed; the cells were reduced in number, flattened and enlarged (Supplementary Figure 8). DOX is a well-known inducer of senescence and in accordance with this, the same phenotypes were observed at low DOX concentrations. In contrast, high concentrations of both LQ-10% and free DOX led to cell death (Supplementary Figure 8).

**Figure 6.**
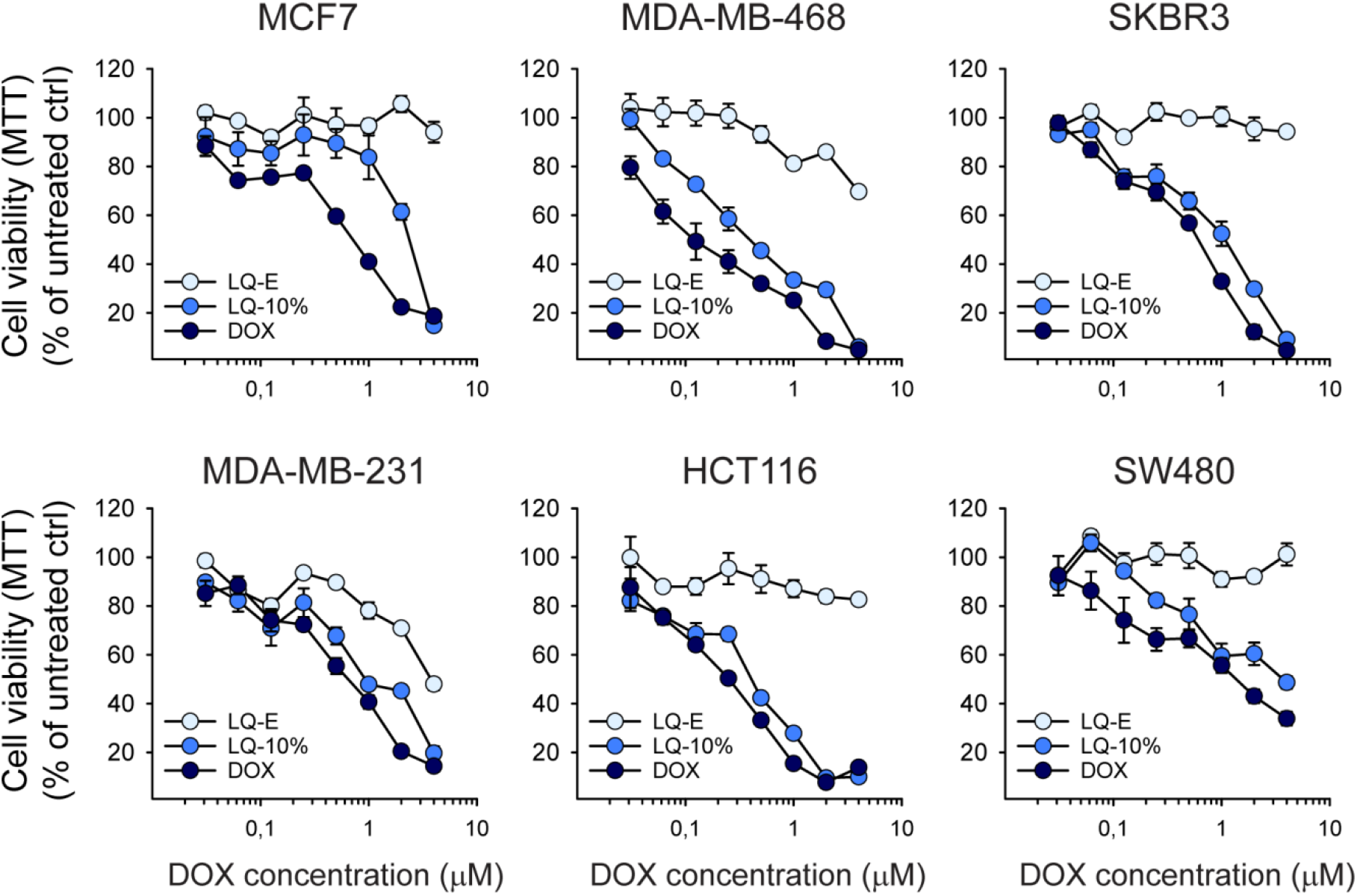
Cytotoxicity of Lipodisqs versus free DOX across a panel of cancer cell lines. The indicated cell lines were treated for 48 h with increasing concentrations of free DOX or with equivalent concentrations of DOX-loaded Lipodisqs (LQ-10%) and the same dilutions of empty Lipodisqs (LQ-E). Cell viability was assessed by the MTT assay. All values were normalised to that of untreated control cells, and the error bars represent standard deviation between triplicates from one representative experiment. Note that the values of the errors are sometimes too low to be detected behind the large circles.

### Biodistribution of Lipodisqs in mice showed a high liver uptake and rapid biliary excretion

To study the biodistribution of Lipodisq NPs, the Cy5 fluorophore was covalently conjugated to either the SMA polymer or to the lipid DLPE (see Supplementary Table 1). Whole body images were obtained using the IVIS^®^ Spectrum scanner, and mice were sacrificed at the given time points such that organs could be harvested and visualised *ex vivo* (Supplementary Figure 9 and 10). The majority of fluorescence from both Lipodisq preparations was recovered in liver (Supplementary Figure 10), which is consistent with previous observations for other *i.v*. injected NPs [31]. However, contrary to that reported for most NPs, only a small fraction of Lipodisq NPs was recovered in spleen. Notably, there seemed to be a higher fluorescence in animals receiving SMA-Cy5 than DLPE-Cy5 (Supplementary Figure 9 and 10). It is not likely that this difference is caused merely by a higher fluorescence of the SMA-Cy5 preparation, since the fluorescence of SMA-Cy5 was found to be only 11% higher than that of DLPE-Cy5 before injection (data not shown). We favour the idea that some DLPE-Cy5 may be released (see below). Furthermore, surprisingly high levels of fluorescence following injections of both these substances were observed in colon already 1 h after injection, indicating a rapid biliary excretion (Supplementary Figure 10). To study the colon accumulation in more detail, the colon was imaged in the subsequent study also after removal of feces, and both feces and empty colon was imaged (Figure 7). The images and the quantitative data obtained clearly demonstrate that the fluorophore is rapidly excreted into feces (Figure 7). Thus, our biodistribution data show a high recovery of Lipodisqs in liver, followed by a rapid biliary excretion into feces of some of the NPs or Cy5-containing substances. We suggest that the liver uptake is due to the ability of these very small NPs (10 nm in diameter, 4.5 nm thickness) to pass the fenaestra of the liver sinusoidal endothelial cells [32], before being endocytosed by hepatocytes [33] and then excreted into bile [34].

**Figure 7.**
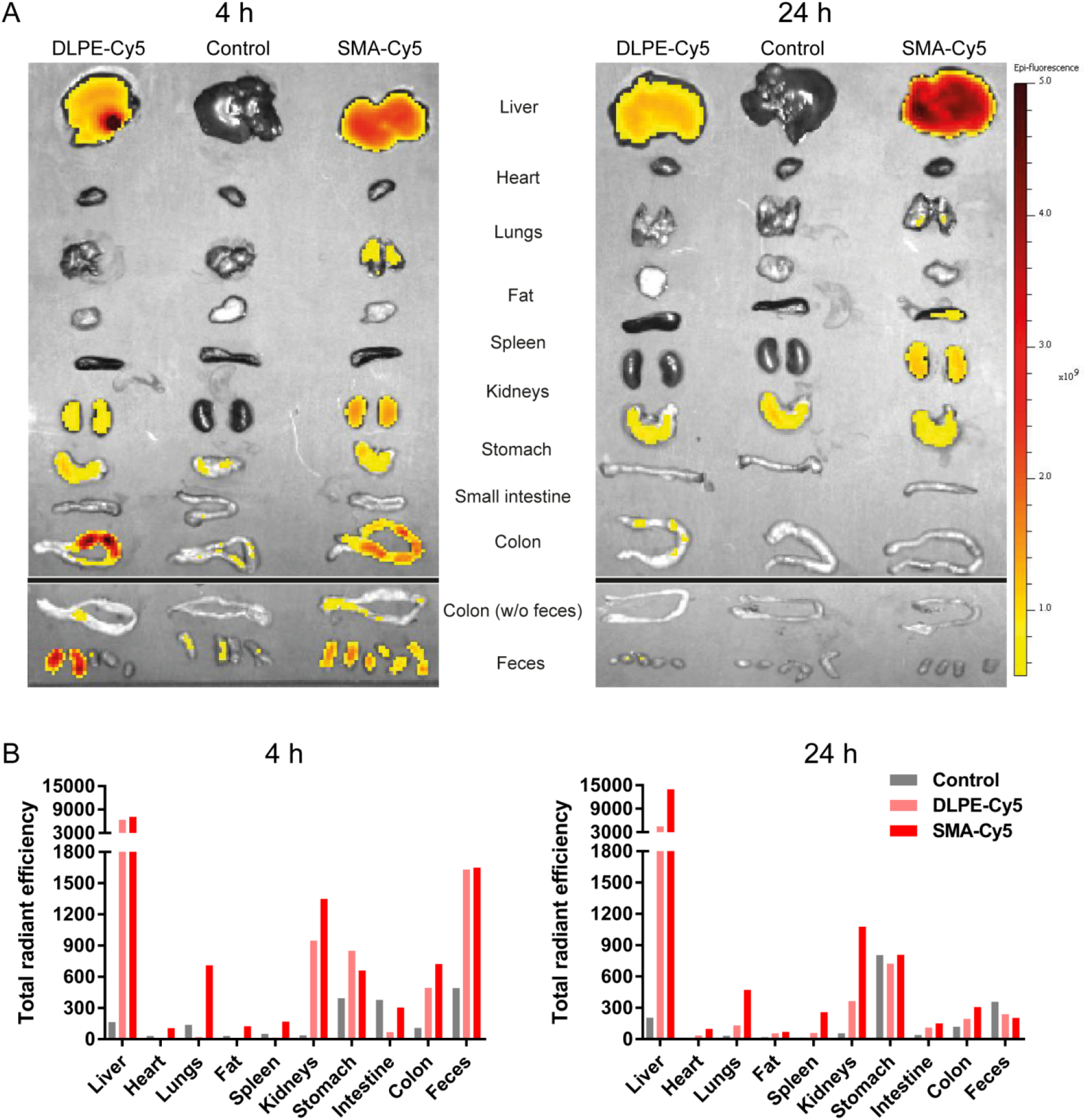
*Ex vivo* fluorescence images of isolated organs after injection of Cy5-labelled Lipodisqs. **A.** Organs obtained 4 and 24 h after injection of SMA-Cy5, DLPE-Cy5 and saline as a control. Colon was imaged both as harvested and after removal of feces. **B.** Quantification of fluorescence obtained from the tissues in A. The data are not corrected for a 11% higher fluorescence of SMA-Cy5 compared to DLPE-Cy5.

By comparing the data obtained 4 and 24 h after injection (Figure 7 and Supplementary Figure 10), it is clear that DLPE-Cy5 is excreted faster than SMA-Cy5. This difference may be due to metabolic release of some of the DLPE-Cy5 lipid from these NPs. In contrast to the other tissues, the fluorescence from SMA-Cy5 strongly increased in liver from 4 to 24 h after injection.

The biodistribution data showing high liver uptake and rapid biliary excretion of Lipodisqs following *i.v*. injection in mice (Figure 7), differ slightly from biodistribution data recently reported for similar NPs [35]. Tanaka *et al*. made SMA-containing partices (8-10 nm) using the phospholipid PC16:0/18:1, and measured biodistribution following *i.v*. injection detected by ^111^In-DTPA bound to PE18:0/18:0. They reported that per g tissue, the spleen contained 31% and 25% of that in liver at 3 and 24 h after injection, respectively. For comparison, the numbers obtained in our study 4 and 24 h after injection was 1.4% and 7.1%, respectively, for the DLPE-Cy5 labelled NPs and 12.6% and 13.6% for the SMA-Cy5 labelled NPs. Furthermore, Tanaka *et al*. [35] reported that the amount of radioactivity in kidneys (per g tissue) was approx. 10% of that in liver at both time points, whereas much higher values were obtained for the Lipodisq NPs, (39% and 28% for the DLPE-Cy5 labelled NPs at 4 and 24 h after injection and 62% and 18% for the SMA-Cy5 labelled NPs). The much higher relative content in kidney than liver for the Lipodisqs are most likely explained by the rapid removal from liver by biliary excretion of these NPs. Tanaka *et al*. did not discuss biliary excretion or the potential presence of their NPs in feces.

Huda *et al*. [36] have also performed biodistribution studies of discoidal NPs formed by using PC16:0/18:1 and MSP1E3D1 protein in place of SMA. NPs of 13 nm were obtained and the biodistribution was evaluated by PET/CT following injection of particles labelled with ^64^Cu-DOTA to lysine groups in the protein. These NPs were reported to permeate deeply into cancer tissue, to show a high accumulation in kidney and a rapid renal excretion [36]. A rapid renal excretion of NPs of 13 nm is unexpected [37]. As these ^64^Cu-labelled NPs released considerable amounts of radioactivity following *in vitro* incubation with mouse plasma at 37 °C for 12 h [36], it is tempting to speculate the at least part of the radioactivity recovered in kidneys was due to release of the radiochemical labelling of the NPs, rather than the excretion of intact NPs.

The biodistribution pattern of Lipodisq NPs reported here, suggests that they are promising candidates for hydrophobic drug delivery to liver. Moreover, the rapid excretion into bile is interesting regarding the possibility to use these NPs for *in vivo* imaging. For a contrast agent to be useful for medical imaging, it is very important that the agent is rapidly excreted to produce a good contrast between the diseased area and the surrounding tissue. In most cases, rapid excretion is essential to obtain a low signal in the surrounding tissue [38]. Most contrast agents for medical imaging are sufficiently small and hydrophilic to be rapidly excreted into urine. This has, however, the drawback that it may be difficult to image diseased areas close to the kidneys or the urinary tract. NPs that are rapidly excreted into bile may therefore be a suitable platform for new contrast agents. In addition, the rapid biliary excretion of Lipodisqs, and also their similarities to HDL [6], suggests that these NPs may be suitable for delivery of drugs and contrast agents to atherosclerotic tissue [39]. Moreover, it may be possible to change or optimalize the pharmacokinetics of these NPs, by coupling PEG to the Lipodisqs (either to the lipids or the SMA polymer) as recently demonstrated for synthetic HDL NPs [40]. In summary, Lipodisqs should be explored further as candidates for the delivery of drugs and contrast agents.

## Abbreviations

DLPE: 1,2-dilauroyl-*sn*-glycero-3-phosphoethanolamine
DLPE-Cy5: Cy5 dye covalently conjugated to 1,2-dilauroyl-*sn*-glycero-3-phosphoethanolamine
DLS: dynamic light scattering
DMPC: 1,2-dimyristoyl-*sn*-glycero-3-phosphocholine
DMPG: 1,2-dimyristoyl-*sn*-glycero-3-phospho-(1’-rac-glycerol)
DOX: Doxorubicin
EPR: enhanced permeability and retention effect
Lipodisq: LQ
NMR: nuclear magnetic resonance
NP: nanoparticle
RT: room temperature
SMA: hydrolysed co-polymer of styrene and maleic anhydride
SMA-Cy5: Cy5 dye covalently conjugated to a thiol derivative of SMA
SMAnh: non-hydrolysed co-polymer of styrene and maleic anhydride
SMAnh-SH: thiol derivative of SMAnh
SMA-SH: thiol derivative of SMA

## Acknowledgements

The work in Oslo was supported by the Research Council of Norway (NANO2021; NANOCAN project no. 228200 and nanoAUTOPHAGY project no. 274574), The Norwegian Cancer Society and Helse Sør-Øst, Norway. We thank the Department for Comparative Medicine for the services they provided. The work in Oxford was supported by the UK EPSRC (research grant number EP/I029516/1), EURAMET (BiOrigin research grant HLT10) and the National Physical Laboratory.

