## Supplementary Material for "Physicochemical characterization, toxicity and *in vivo* biodistribution studies of a discoidal, lipid-based drug delivery vehicle: Lipodisq nanoparticles containing doxorubicin"

Supplementary Figures and Tables

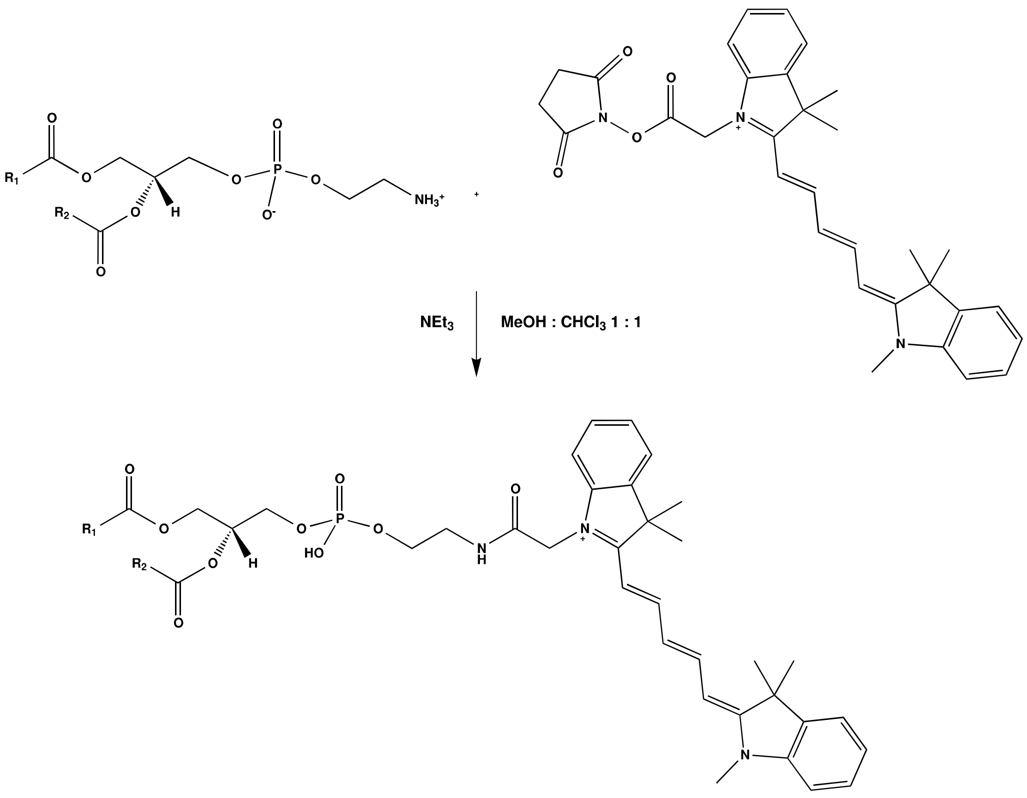


**Supplementary Figure 1. Conjugation scheme for DLPE lipid to the Cy5-NHS ester.** The synthesis was performed using 1,2-dilauroyl-*sn*-glycero-3-phosphoethanolamine (DLPE; PE 12:0/12:0). The figure shows, however, the reaction for any PE species. The conjugation reaction was performed in a 1:1 (v/v) mixture of methanol (MeOH) and chloroform ($CHCl_3$) in the presence of *N*,*N*-diethylethanamine ($NEt_3$).

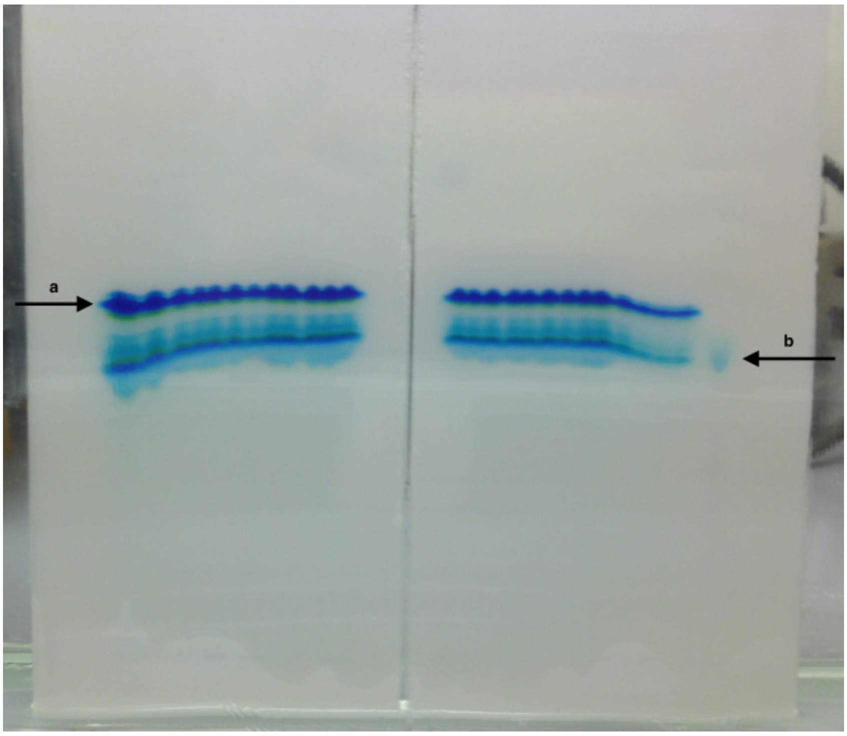


**Supplementary Figure 2. Preparative TLC for separation of DLPE-Cy5 from Cy5-NHS dye conjugate.** The top band (a) is the DLPE-Cy5 whereas the bottom band (b) is unreacted Cy5-NHS dye.

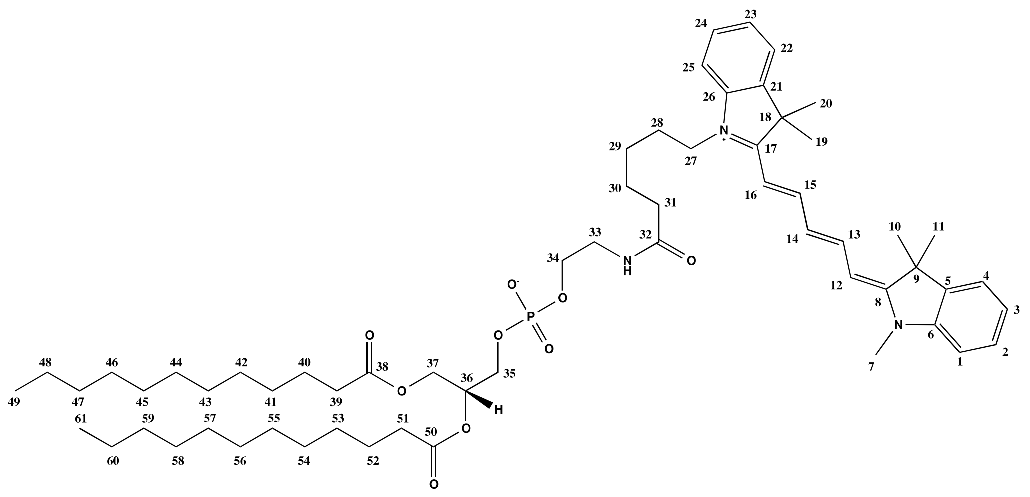

**Supplementary Figure 3. NMR and mass spectrometry analyses of synthesized DLPE-Cy5.** The following data show that DLPE-Cy5 was obtained. NMR:
δH (500 MHz, CDCl3) = 0.69 - 0.89 (H49 + H61, m, 6 H), 1.08 - 1.27 (H40 - H47 + H52 - H59, m, 30 H), 1.32 (H48 + H60, m, 4 H), 1.36 (H27 + H31 + H39 + H51, m, 8 H), 1.50 (H10 + H11+ H19 + H20, s, 12 H), 1.69 - 1.74 (H29, m, 2 H), 2.09 - 2.29 (H28 + H30, m, 4 H), 3.36 - 3.50 (H33, m, 2 H), 4.00 (H35, m, 2 H), 4.08 - 4.17 (H37, m, 2 H), 4.15 - 4.33 (H34, m, 2H), 4.58 - 4.66 (H7, s, 3 H), 5.01 (H36, m, 1 H), 6.42 - 6.44 (H12 + H16, m, 2 H), 6.58 -6.61 (H14, m, 1 H), 6.99 - 7.07 (H1 + H25, m, 2 H), 7.10 - 7.16 (H4 + H22, m, 2 H), 7.24 - 7.37 (H2 + H3 + H23 + H24, m, 4 H), 7.60 - 7.74 (H13 + H15, m, 2 H); δC (126 MHz, CDCl3) = 14.08 (C49 + C61), 22.72 (C10 + C19), 23.76 (C27 + C31 + C39 + C51), 25.80 (C29), 28.11 - 30.37 (C40 - C48 + C53 - C60), 31.95 (C11 + C20), 36.12 (C28 + C30), 41.75 (C33), 44.41 (C35), 62.92 (C34), 63.38 - 63.50 (C37), 70.69 - 70.76 (C36), 104.26 (C12 + C16), 105.39 (C14), 110.15 (C1 + C25), 111.18 (C7), 124.67 - 125.24 (C4 + C22), 126.39 - 127.32 (C9 + C18), 128.64 - 132.47 (C2 + C3 + C23 + C24), 140.66 - 141.12 (C8 + C17), 142.02 (C6 + C26), 143.02 (C5 + C21), 152.28 - 153.18 (C13 + C15),167.79 (C32), 172.12 - 173.47 (C38 + C52); $v_{max}$ 1734, 1717. Mass spectrometry analysis: DLPE-Cy5 has the chemical composition $C_{61}H_{94}N_3O_9P$; the theoretical *m/z* value for the sodium adduct, i.e. $[M+Na]^+$, is 1044.68004; found 1044.67879.

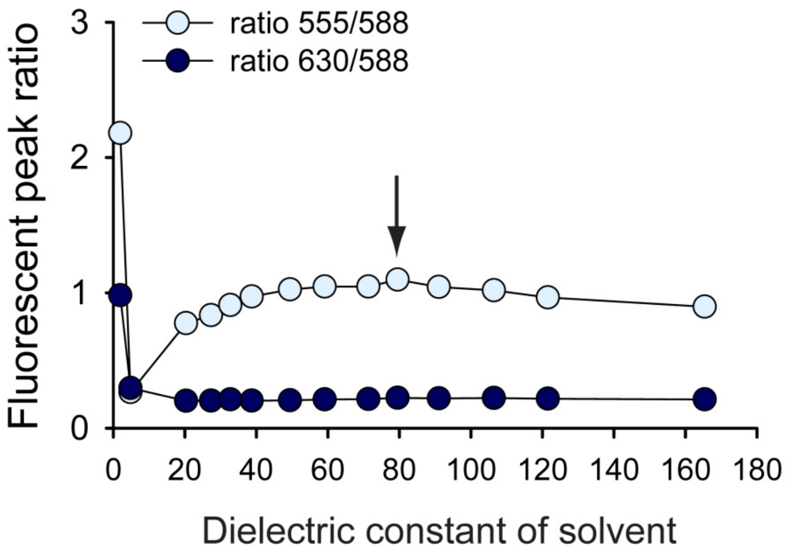


**Supplementary Figure 4. Fluorescence peak ratios of DOX in solvents of different dielectric constant.** DOX was dissolved to a concentration of 30 µM in each solvent, except the two with the lowest dielectric constants (hexane and chloroform) which were saturated with the maximum concentration of DOX, but which was lower than 30 µM. The black arrow overlaid on the graph indicates water as the solvent.

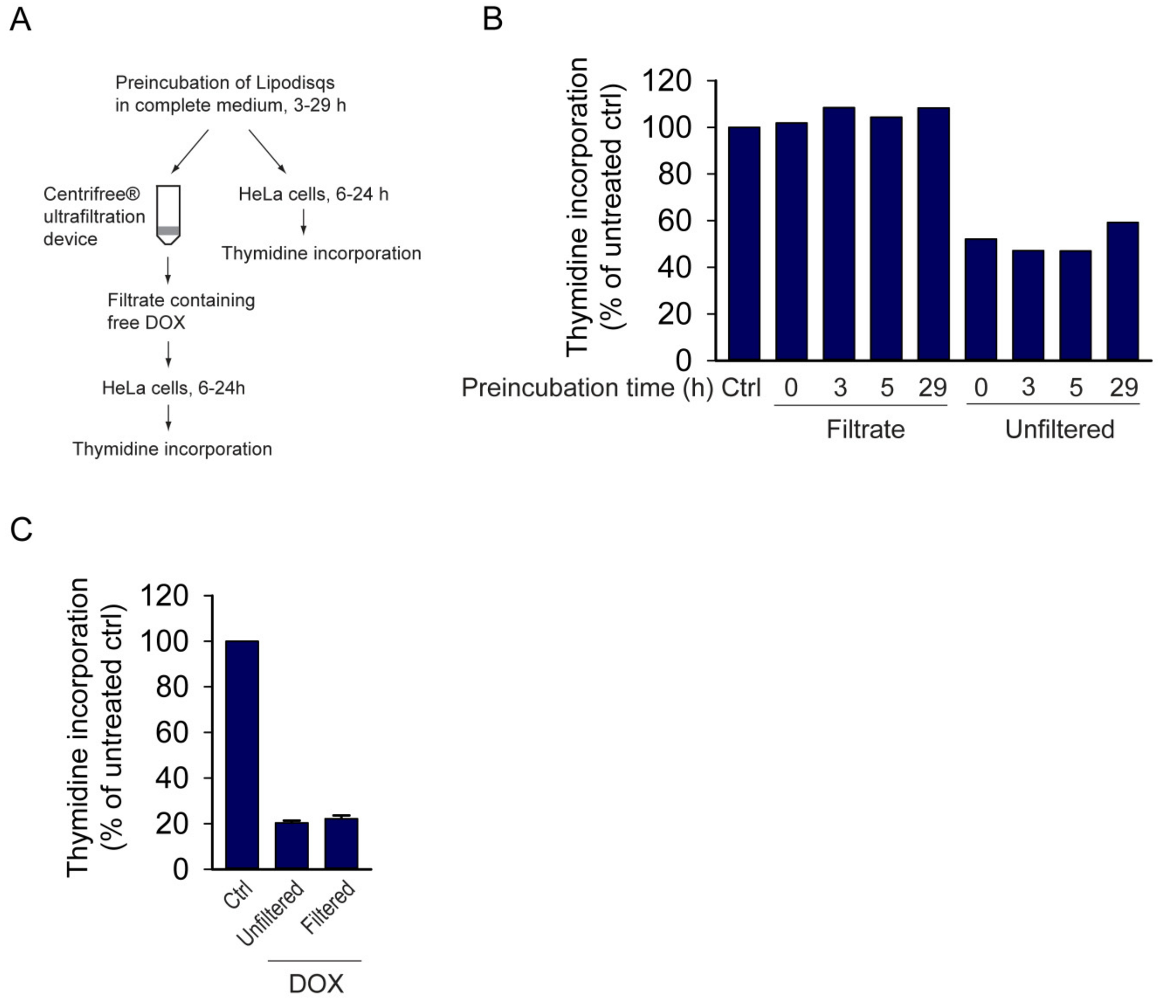


**Supplementary Figure 5. Lipodisqs do not release DOX in complete growth medium in the absence of cells. A.** Workflow for the use of Centrifree® Ultrafiltration device for crude determination of DOX release from Lipodisqs. **B.** LQ-5% (1:100) was incubated at 37 °C for 3, 5 or 29 h in complete growth medium. Then half of the Lipodisq solution was passed through a Centrifree® Ultrafiltration device. The filtrate or the unfiltered Lipodisqs were diluted further 1:2 in complete medium and added to HeLa cells and incubated for 16 h before thymidine incorporation was measured. **C.** HeLa cells were treated with free DOX (340 nM) for 24 h before measuring thymidine incorporation. Free DOX exerts the same degree of cytotoxicity whether the solution is filtered or not, showing that DOX is able to pass freely through the filter.

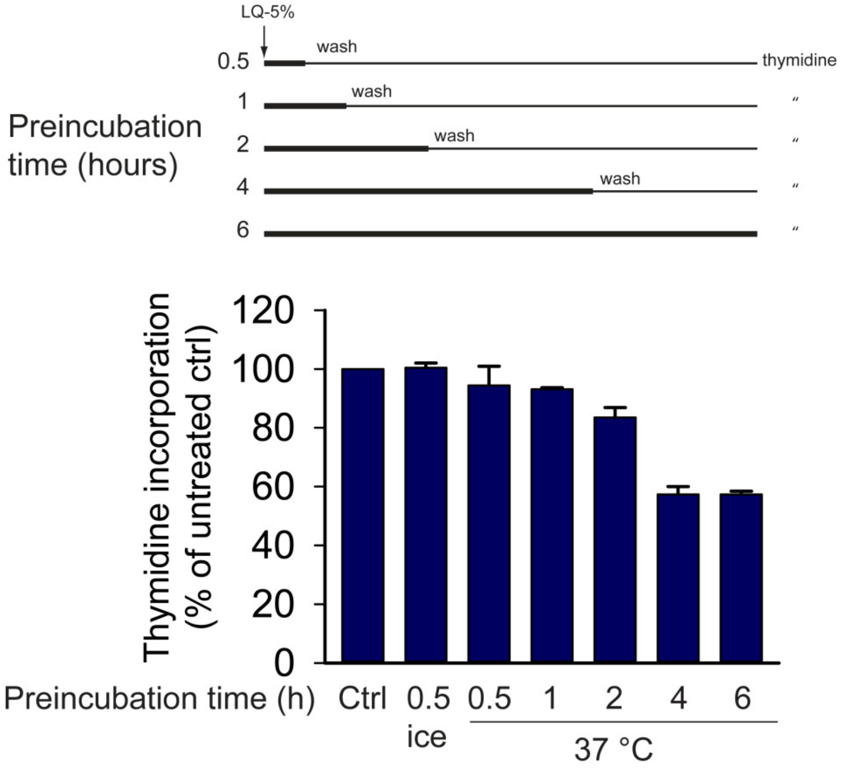


**Supplementary Figure 6. Four hours of Lipodisq uptake is sufficient for maximal cytotoxic effect.** HeLa cells were treated with Lipodisqs (LQ-10% 1:100) for 30 min on ice or for 0.5-6 h at 37 °C, before unbound Lipodisqs were removed by washing. Thereafter the incubation was continued, and 6 h after Lipodisq addition, thymidine incorporation was assessed by incubation with [$^3$H]thymidine for 30 min. The data were normalised to untreated cells, and the error bars show deviation from mean from duplicates.

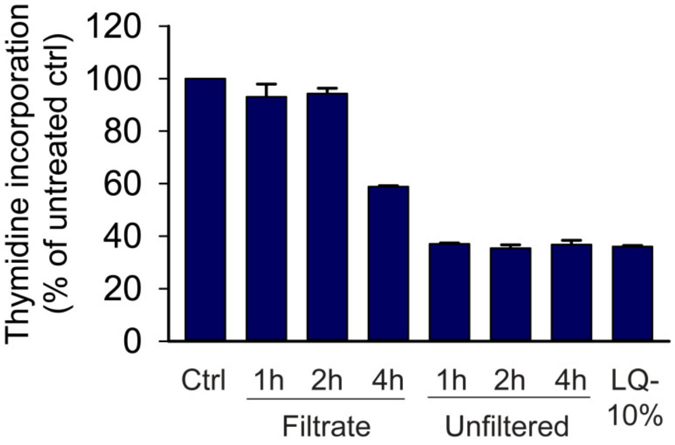


**Supplementary Figure 7. The release of DOX at low pH is not a spontaneous process, but rather takes time.** LQ-10% (1:25) was incubated in PBS pH 5.0 at 37 °C for 1, 2 or 4 h. Then the solution was filtered through a Centrifree® spin column to separate free DOX from Lipodisq-bound drug. The filtrate or the unfiltered solution was diluted 1:40 in complete medium and added to HeLa cells. The incubation was continued for 24 h before the level of thymidine incorporation was determined. The error bars show deviation from mean from duplicates.

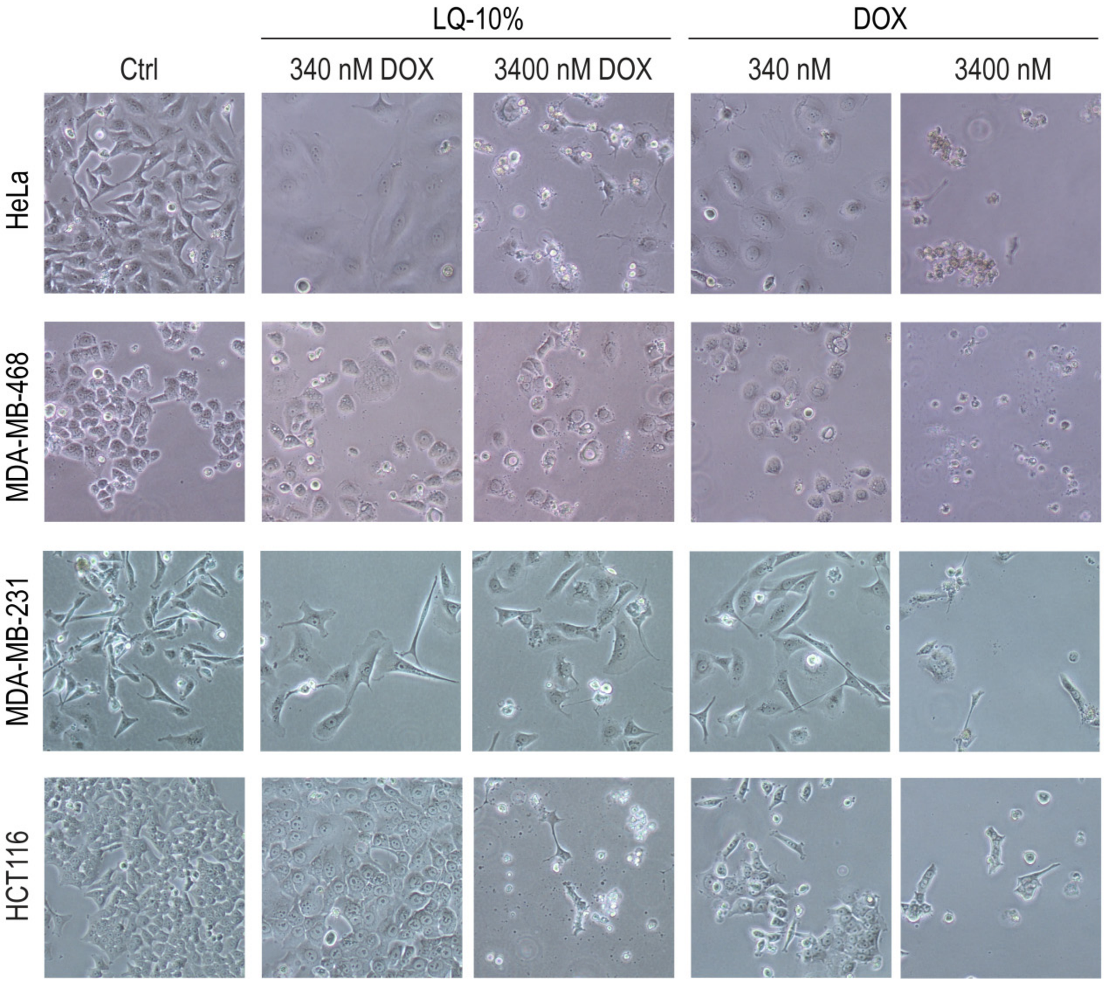


**Supplementary Figure 8. The Lipodisq NPs LQ-10% induce senescence or cell death in a concentration-dependent manner.** The indicated cell lines were treated with 340 or 3400 nM free DOX or LQ-10% containing equivalent DOX concentrations. Brightfield images were aquired after 48 h. At 340 nM DOX, phenotypes characteristic of senescence, such as cell flattening and enlargement, were observed in all cell types, whereas 3400 nM of DOX induced cell death.

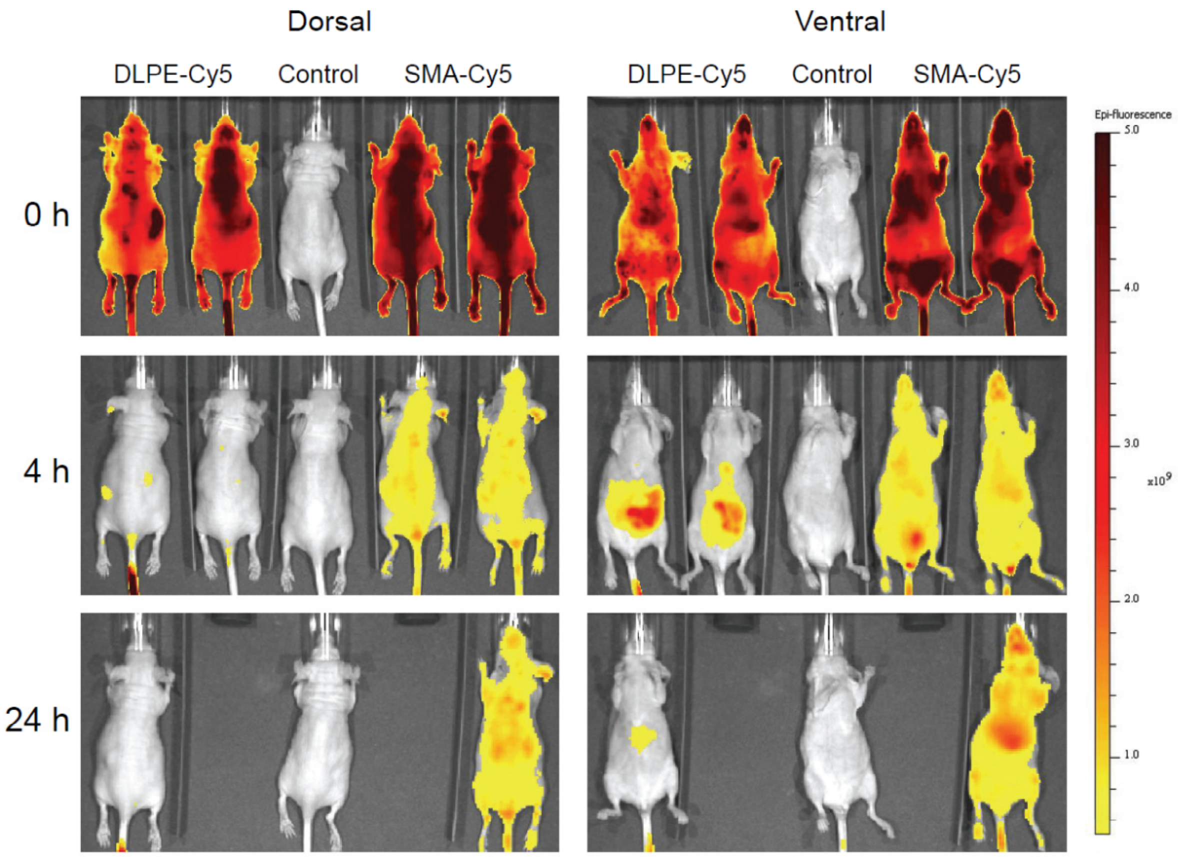


**Supplementary Figure 9. Whole body fluorescence images.** Images with dorsal and ventral views were obtained immediately after injection (0 h) as well as 4 and 24 h after injection of DLPE-Cy5, SMA-Cy5 or saline as control. The images are not corrected for an 11% higher fluorescence of SMA-Cy5 compared to DLPE-Cy5.

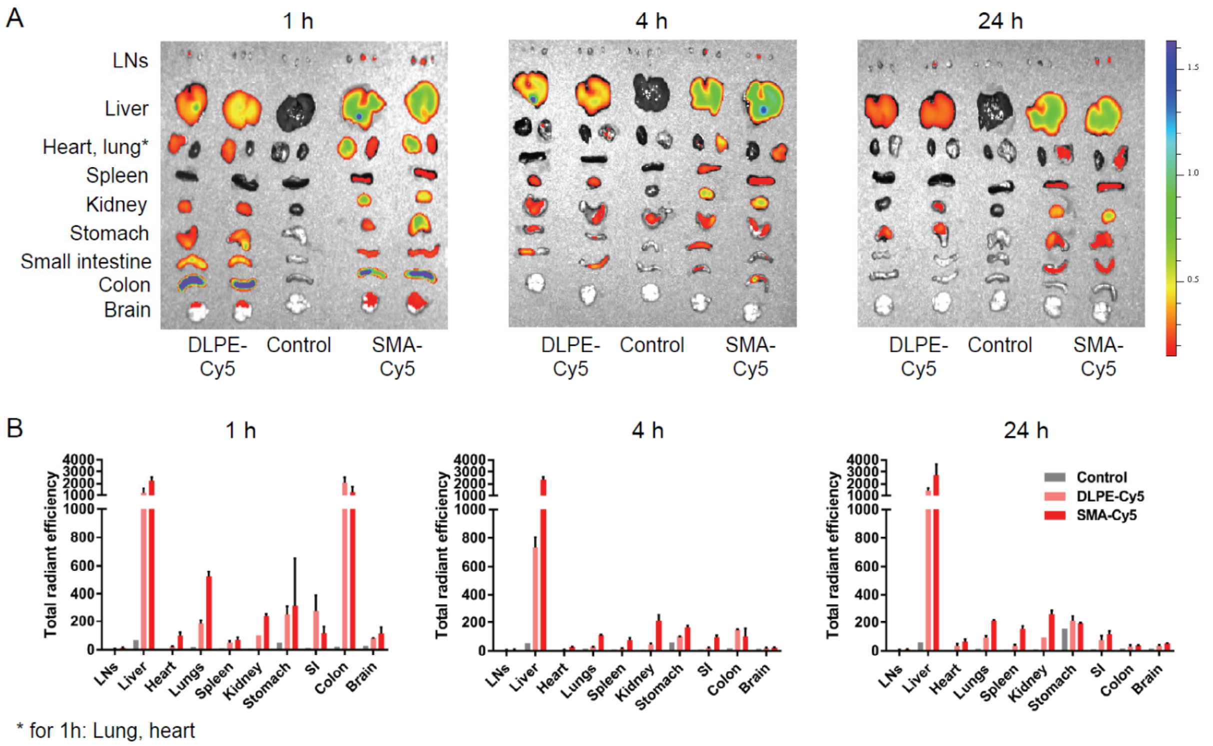


**Supplementary Figure 10. *Ex vivo* fluorescence images of isolated organs after injection of Cy5-labelled Lipodisqs**. **A.** Organs obtained 1, 4 or 24 h after injection of DLPE-Cy5, SMA-Cy5 or saline as control. LNs = lymph nodes. **B.** Quantification of fluorescence obtained from the tissues in A.

**Supplementary Table 1. The composition of Lipodisq batches produced for the different applications.**

| Application | Lipodisq name | Lipid | DOX (molar %) | Dye |
|---|---|---|---|---|
| Physiochemical characterization | DMPC Lipodisqs | DMPC | - | - |
| | DMPG Lipodisqs | DMPG | - | - |
| Cellular uptake and cytotoxicity | LQ-E | DMPC 2.2 mg/ml (3.2 mM) Cy5-DLPE 1% | - | Cy5 conjugated to DLPE |
| | LQ-1% | DMPC 2.2 mg/ml Cy5-DLPE 1% | 1% DOX | Cy5 conjugated to DLPE |
| | LQ-5% | DMPC 2.2 mg/ml Cy5-DLPE 1% | 5% DOX | Cy5 conjugated to DLPE |
| | LQ-10% | DMPC 2.2 mg/ml Cy5-DLPE 1% | 10% DOX (360 µM) | Cy5 conjugated to DLPE |
| Biodistribution in mice | Lipodisq DLPE-Cy5 | DMPC 10 mg/ml, Cy5-DLPE 1% | - | Cy5 conjugated to DLPE |
| | Lipodisq SMA-Cy5 | DMPC 10 mg/ml | - | Cy5 conjugated to SMA |

**Supplementary Table 2. Dielectric constant of solvents and mixtures in which DOX was dissolved.**

| Solvent | Dielectric constant |
|---|---|
| Hexane | 1.9 |
| Chloroform | 4.8 |
| Propan-2-ol | 20.3 |
| EtOH | 27.3 |
| MeOH | 32.7 |
| MeOH + $H_2O$ (4:1) | 38.7 |
| MeOH + $H_2O$ (3:7) | 49.4 |
| MeOH + $H_2O$ (1:1) | 59.0 |
| MeOH + $H_2O$ (1:9) | 71.3 |
| $H_2O$ | 79.5 |
| NMF + $H_2O$ (3:7) | 91.0 |
| NMF + $H_2O$ (1:1) | 106.4 |
| NMF + $H_2O$ (7:3) | 121.5 |
| NMF + $H_2O$ (95:5) | 165.4 |
